# New neurons in old brains: A cautionary tale for the analysis of neurogenesis in post-mortem tissue

**DOI:** 10.1101/2021.11.12.468443

**Authors:** Dylan J. Terstege, Kwaku Addo-Osafo, G. Campbell Teskey, Jonathan R. Epp

## Abstract

Adult neurogenesis has primarily been examined in two key regions in the mammalian brain, the subgranular zone of the hippocampus and the subventricular zone. The proliferation and integration of newly generated neurons has been observed widely in adult mammalian species including the human hippocampus. Recent high-profile studies have suggested however, that this process is considerably reduced in humans, occurring in children but declining rapidly and nearly completely in the adult brain. In comparison, rodent studies also show age-related decline but a greater degree of proliferation of new neurons in adult animals. Here, we examine whether differences in tissue fixation, rather than biological difference in human versus rodent studies might account for the diminished levels of neurogenesis sometimes observed in the human brain. To do so we analyzed neurogenesis in the hippocampus of rats that were either perfusion-fixed or the brains extracted and immersion-fixed at various post-mortem intervals. We observed an interaction between animal age and the time delay between death and tissue fixation. While similar levels of neurogenesis were observed in young rats regardless of fixation, older rats had significantly fewer labeled neurons when fixation was not immediate. Furthermore, the morphological detail of the labeled neurons was significantly reduced in the delayed fixation conditions at all ages. This study highlights critical concerns that must be considered when using post-mortem tissue to quantify adult neurogenesis.

---

Adult neurogenesis occurs in at least two regions of the mammalian brain, the dentate gyrus of the hippocampus (HPC) and the subventricular zone. Among mammalian species, neurogenesis has been most widely studied in rodents. There is a well-documented age-dependent decline in adult neurogenesis in rodents but, the presence of new neurons remains detectable throughout adult life. With few exceptions[1], adult neurogenesis has been observed across mammalian species[2,3] suggesting that it is highly conserved. However, in humans, there is some disagreement over the extent and timecourse of postnatal neurogenesis.

Adult neurogenesis in the human brain was first observed in cancer patients administered Bromodeoxyuridine (BrdU) to track tumor growth[4]. BrdU, an exogenous thymidine analogue is incorporated into dividing cells during DNA synthesis. Confirmation of neuronal phenotype is then provided via double immunohistochemistry with neuron specific markers such as NeuN. Post-mortem analysis of brains from these individuals indicated continued neurogenesis in adulthood as determined by the presence of BrdU/NeuN double labeled cells in the HPC. Numerous studies have confirmed these findings using several methods to identify new neurons (for review see[5]). The majority of studies of neurogenesis in humans used immunohistochemical labeling of endogenous markers of proliferation[6,7] such as Ki67, and/or immature neurons[8]. The most commonly used marker for neurogenesis, a protein called doublecortin (DCX), is highly enriched in immature neurons[9]. In most cases, adult generated neurons have been identified across the lifespan from 0 – 100 years of age[10] with an age related decrease similar to other species[10,11]. The methods used to identify new neurons in the human brain have some caveats such as potential damage/repair induced uptake of BrdU[12].

However, the cumulative evidence from multiple approaches strongly suggests the presence of adult neurogenesis in the human HPC. Despite this, several recent and controversial papers reported the absence of adult generated neurons in humans although they did observe new neurons in juveniles [13–15]. These studies concluded that if neurogenesis occurs at all in the adult human brain it is a very rare event. With respect to the many other studies that documented neurogenesis in the adult human HPC, it was suggested that these positive findings may actually represent non-specific or non-neuronal labeling rather than adult neurogenesis[14].

One potential concern when analyzing adult neurogenesis in humans relates to the collection of post-mortem tissue and the degree to which the tissue degrades prior to complete fixation. Nonhuman animal studies allow for highly controlled tissue collection that normally involves perfusion fixation, during which fixative is introduced through the vasculature at the time of sacrifice. As a result, the interior of the brain is quickly exposed to the fixative which prevents autolysis and allows for relatively rapid, and consistent fixation. In most cases, human tissue is collected post-mortem and the time-window between death and tissue collection is highly variable. The longer it takes to initiate fixation the greater the degree of tissue/protein degradation that may occur[16–18]. In addition, penetration of the fixative (and time for subsequent protein cross-linking) when an immersion fixation technique is employed, is not instantaneous. Depending on tissue size it can take hours to days for the interior aspects of a tissue block to be exposed and fixed by the commonly used aldehyde fixatives even if the post-mortem tissue was collected rapidly[19]. Differences in fixation methods present a challenge in comparing animal studies of adult neurogenesis with human studies. The goal of this study was to determine in rats the effects of post-mortem delays in tissue collection, on the ability to detect adult hippocampal neurogenesis.

We examined brains from 4-month or 9-month-old male adult Sprague Dawley rats (Charles River Laboratories, Kingston, NY, USA) that were transcardially perfused to age matched rats that were killed, and their brains collected at 0-, 6-, or 12-hours post-mortem. Rats were chosen because their brain volume (∼mm3) is similar to the minimum human tissue block sizes used previously[14]. Rats in the perfused group were deeply anesthetized and perfused with 60 ml of PBS followed by 120 ml of 4% formaldehyde. The brains were extracted, and immersion fixed in the same fixative for 48 hours. For the delay fixation conditions, rats were given a lethal overdose of isoflurane (5% ISO, until death) and the brains were extracted following a delay of either 6 or 12 hours (Fig. 1a). Brains were placed in 4% formaldehyde for 48 hours. After fixation brains were transferred to 30% sucrose solution and were sectioned on a Cryostat (Leica CM1950) at a thickness of 40 µm. Tissue series (1/12) were stored at -20 °C in antifreeze.

**Figure 1.**
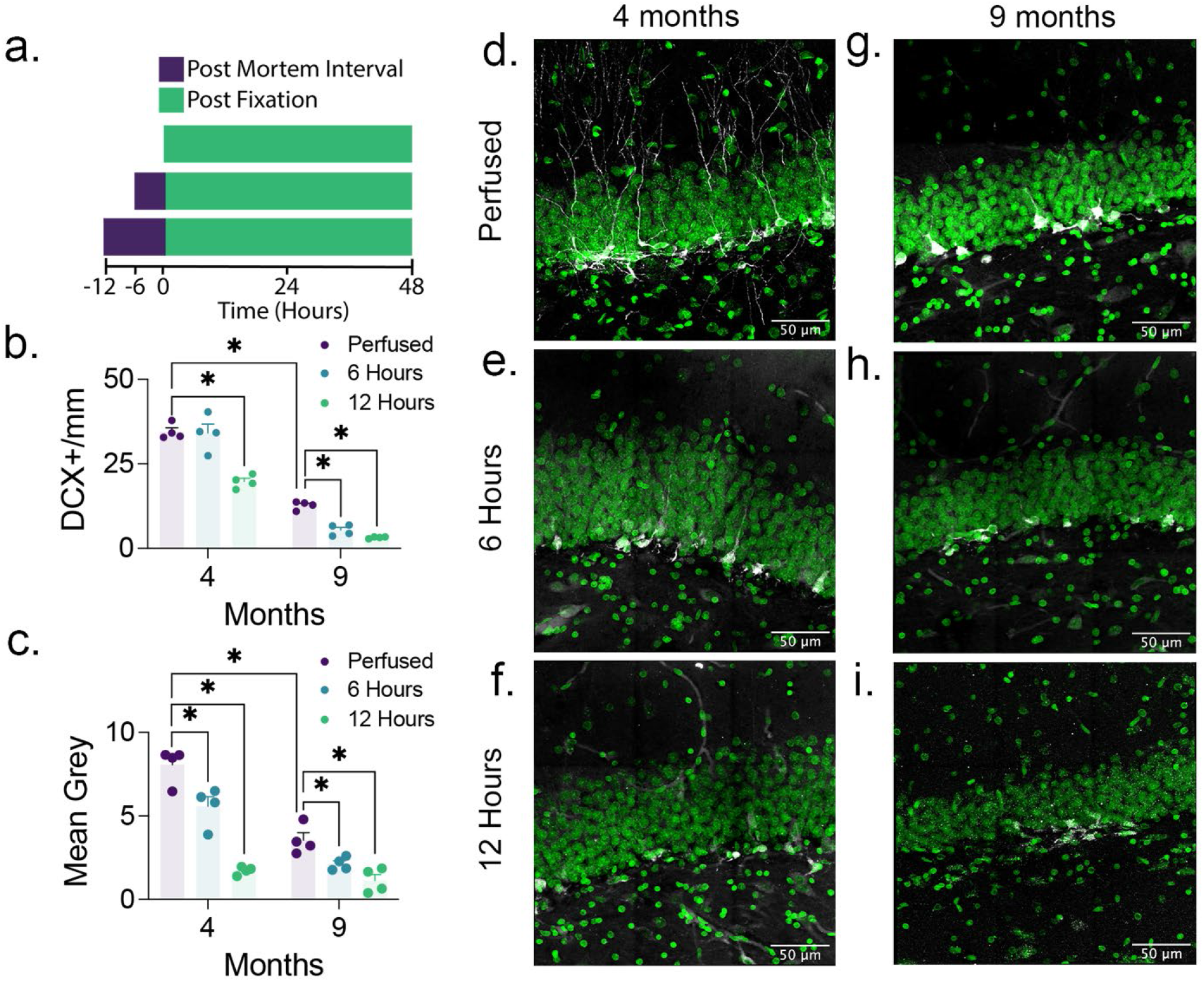
Doublecortin labeling is reduced with increased post-mortem interval. (**a**) Following post-mortem intervals of 0, 6, or 12 hours, brains from 4- and 9-month-old rats (*n* = 4 were fixed for 48 hours. The effects of age and post-mortem interval interacted significantly in influencing the (**b**) density of doublecortin expression (Two-Factor ANOVA; F_2,18_ = 10.97, *p* = 0.0008) and the (**c**) optical density of doublecortin labelling in the granule cell layer and subgranular zone (Two-Factor ANOVA; F_2,18_ = 12.34, *p* = 0.0004). Representative images of doublecortin expression in 4-month-old (**d**-**f**) and 9-month-old (**g**-**i**) rats with post-mortem intervals of 0 (**d**,**g**), 6 (**e**,**h**), and 12 (**f**,**i**) hours. Data are shown as the mean ± SEM. ^*^*P* < 0.05.

DCX labeling was performed as previously described[20] using a 1:250 dilution of primary antibody (Cell Signalling 4601S Rabbit anti-DCX) followed by a 1:500 dilution of donkey anti-rabbit Alexa Fluor 488 labeled secondary antibody (Jackson Immuno Research Laboratories AB_2338072). Quantification of labeled cells was performed on an Olympus BX63 fluorescent microscope at 60x magnification. Labeled cells were counted through the entire extent of the dentate gyrus and were included only if the cell body could be clearly identified and was located in the granule cell layer or subgranular zone. Areas were measured by tracing the DAPI labeled outline of the granule cell layer. Slides were coded so that quantification was performed blind to treatment condition.

The optical density of DCX labelling in the granule cell layer and subgranular zone was performed on an Olympus FV3000 confocal microscope using a 10x objective (0.4 NA). Three images, each from different tissue sections, were collected at a z-spacing of 3.96 μm.

Illumination settings and z-range were consistent across all images. Background mean grey pixel intensity values of the z-projected DCX images were recorded from the outer molecular cell layer using ImageJ. These values were subtracted from mean pixel intensity recorded in the upper blade of the dentate gyrus.

We predicted that post-mortem protein degradation might influence the detection of adult generated neurons and, due to a higher level of protein expression in younger brains, might have a greater influence on detection in older brains. Our results confirm this prediction (Fig. 1b). We found a significant age by fixation interaction effect. Post-hoc analysis of the data using Tukey’s tests indicated the expected reduction in DCX between 4 months and 9 months in perfused rats. Importantly, there was a significant impact of the post-mortem interval on our ability to detect DCX. At the shortest delay of 6 hours there was no significant difference between perfused and non-perfused 4-month-old rats. However, in 9-month-old rats there was a significant decrease in DCX in non-perfused rats. At 12 hours there were significant decreases in DCX in both 4- and 9- month-old rats but, overall, the decrease was considerably greater in the older rats.

We also examined the appearance of DCX labeled cells in each condition (Fig. 2). Perfusion fixation resulted in labeled cells with clear morphology and consistent labeling, while the delayed fixation resulted in poor morphological detail. Dendrites, normally visible following perfusion were less visible, and the cytoplasmic labeling often appeared weak and speckled.

**Figure 2.**
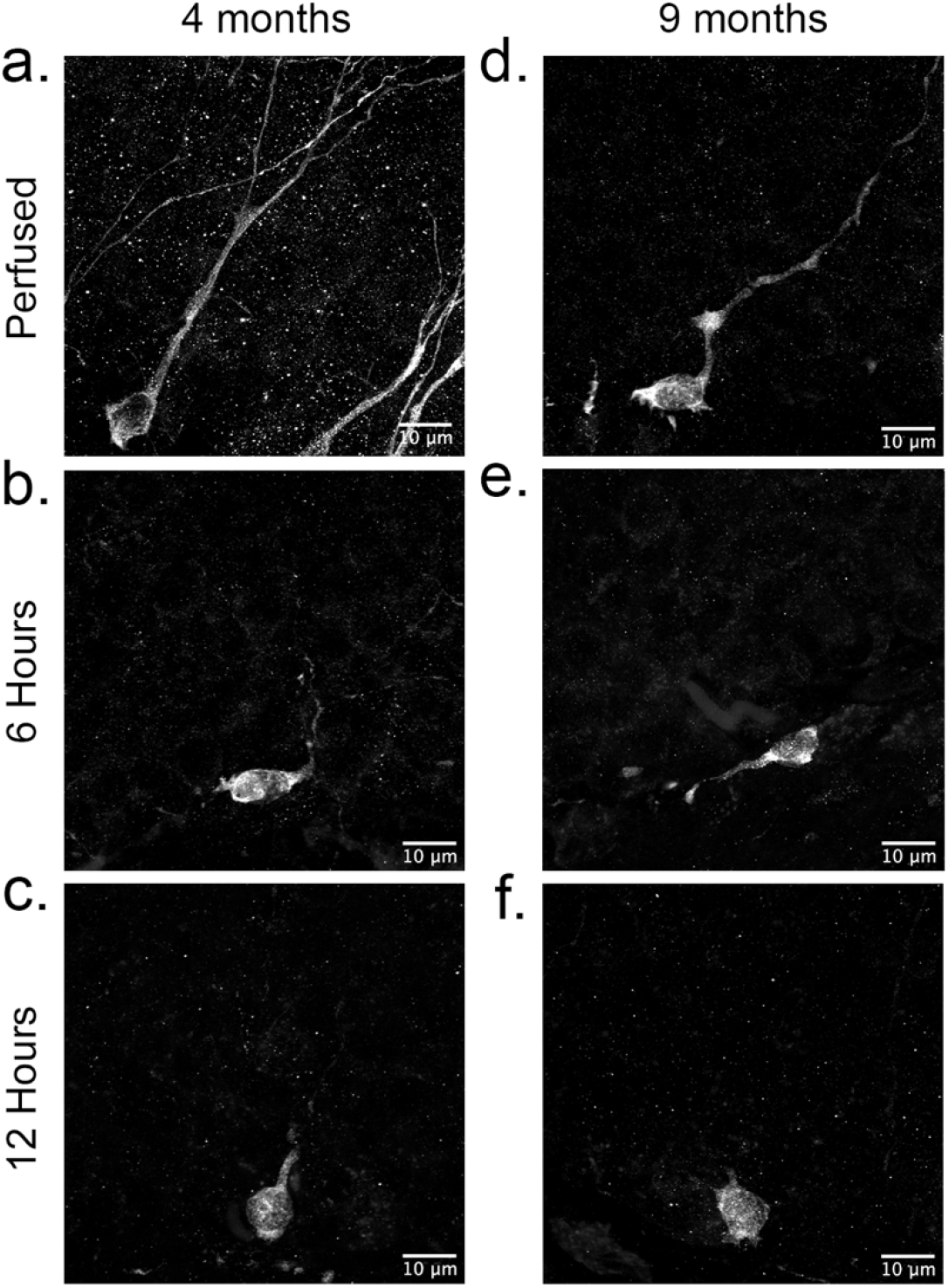
Dendritic morphology in doublecortin-labelled neurons is reduced with increased post-mortem time. Representative images of doublecortin in 4- (**a**-**c**) and 9-month-old (**d**-**f**) rats with post-mortem intervals of 0 (**a**,**d**), 6 (**b**,**e**), and 12 (**c**,**f**) hours prior to fixation.

Thus, identifying the new neurons based on the morphology of the labeling itself becomes increasingly difficult as fixation is compromised. This was further supported by decreased optical density of DCX staining in the combined granule cell layer and subgranular zone of the dentate gyrus (Fig. 1c).

In conclusion, our results demonstrate that age related changes in adult neurogenesis in post-mortem collected tissue must be interpreted with caution as there is potential for misinterpretation of decreased or absent neurogenesis in older subjects. This is especially important if younger tissue is to be used as a positive control. We suspect that the interaction between age and fixation is likely driven by lower levels of DCX protein expression in older animals that are detectable with optimized fixation but more rapidly fall below detection limits as the protein begins to degrade post-mortem. This may occur due to protein loss or conformational changes that prevent antibody binding. The extent to which such an effect may occur with other proteins is unclear but should be considered in future studies. Our current approach may provide a strategy to determine age-related stability of different proteins prior to post-mortem analysis.

## Acknowledgments

Funding for this study was provided by an NSERC Discovery Grant (RGPIN-2018-05135) to JRE and the Canadian Institutes for Health Research (MOP-130495) to GCT. DJT received a fellowship from the Canadian Open Neuroscience Platform. KA received a Queen Elizabeth II fellowship.

